# Higher hyperpolarization activated current (I_h_) in a subpopulation of interneurons in stratum oriens of area CA1 in the hippocampus of Fragile X mice

**DOI:** 10.1101/2022.12.20.521268

**Authors:** Lauren T. Hewitt, Darrin H. Brager

## Abstract

Fragile X syndrome is the most common inherited form of intellectual disability and the leading monogenetic cause of autism. Studies in mouse models of autism spectrum disorders, including the Fmr1 knockout (FX) mouse, suggest that abnormal inhibition in hippocampal circuits contributes to behavioral phenotypes. We and others previously identified changes in multiple voltage-gated ion channels in hippocampal excitatory pyramidal neurons in FX mice. Whether the intrinsic properties of hippocampal inhibitory interneurons are altered in FX remains largely unknown. We made whole-cell current clamp recordings from three types of interneurons in stratum oriens of the hippocampus: fast-spiking (FS) cells and two classes of low threshold spiking cells, oriens-lacunosum moleculare (OLM) and low-threshold high I_h_ (LTH) neurons. We found that in FX mice, input resistance and action potential firing frequency were lower in LTH, but not FS or OLM, interneurons compared to wild type. Bath application of the h-channel blocker ZD7288 had a larger effect on input resistance in FX LTH cells and normalized input resistance between wild type and FX LTH cells, suggesting a greater contribution of I_h_ in FX LTH cells. In agreement, voltage clamp recordings showed that I_h_ was higher in FX LTH cells compared to wild type. Our results suggest that the intrinsic excitability of LTH inhibitory interneurons may contribute to altered inhibition in the hippocampus of FX mice.

**Significance statement:** Changes in the balance between excitation and inhibition underlie many neurodevelopmental disorders. Changes in several voltage-gated ion channels, which contribute to altered intrinsic excitability, were previously identified in excitatory hippocampal neurons in the Fmr1 mouse model of Fragile X syndrome. In this paper we use physiological and biochemical approaches to investigate the intrinsic excitability of inhibitory interneurons in hippocampal area CA1 of the Fragile X mouse. We found that higher I_h_ lowers the intrinsic excitability of one specific type of interneuron. This study highlights how changes to voltage-gated ion channels in specific neuronal populations may contribute to the altered excitatory/inhibitory balance in Fragile X syndrome.

## Introduction

Fragile X Syndrome (FXS) is the primary monogenetic cause of autism and the most common form of inherited intellectual disability. Patients with FXS exhibit epilepsy, sensory hypersensitivity, and anxiety among other cognitive impairments (Hagerman and Hagerman, 2002; Kaufmann et al., 2004; Hall et al., 2008; Hagerman et al., 2009). One working hypothesis is that excitatory/inhibitory (E/I) balance, which is essential for normal brain circuit function, is disrupted in FXS. The hippocampus contains a diverse population of inhibitory interneurons that form distinct circuits that control the precise timing of excitatory activity in the hippocampus (Cobb et al., 1995; Freund and Buzsáki, 1996; Klausberger and Somogyi, 2008; DeFelipe et al., 2013). Given this diversity of interneuron functions, changes in even a single population of interneurons, could skew the balance between excitation and inhibition (Rubenstein and Merzenich, 2003; Belmonte and Bourgeron, 2006; Chao et al., 2010; Paluszkiewicz et al., 2011).

FXS is characterized by the transcriptional silencing of the *Fmr1* gene (Fu et al., 1991) leading to a loss of Fragile X Messenger Ribonucleoprotein 1 (formerly known as Fragile X Mental Retardation Protein, FMRP) ((Bell et al., 1991), FMR1 gene ID: 2332, NCBI, 2022). FMRP is highly expressed in the hippocampus in excitatory pyramidal neurons (Feng et al., 1997); however, there are currently no studies that report the co-expression of FMRP within specific subtypes of inhibitory interneurons in the CA1 hippocampus. FMRP regulates the function and expression of numerous voltage-gated ion channels (Brown et al., 2010; Strumbos et al., 2010; Deng et al., 2013). Within the hippocampus, changes in voltage-gated Na+, K+, and h-channels, that alter intrinsic excitability, were identified in excitatory pyramidal neurons in Fmr1 KO mice (Brager et al., 2012; Routh et al., 2013; Contractor et al., 2015; Deng and Klyachko, 2016; Deng et al., 2019; Brandalise et al., 2020). Whether there are changes in hippocampal inhibitory interneurons in FX mice remain unknown.

Within area CA1 of the hippocampus, stratum oriens interneurons can be divided into two canonical interneuron types based on action potential firing: fast spiking (FS) and low threshold spiking (LTS). FS neurons exert strong, feedforward inhibition onto CA1 pyramidal cell bodies while LTS neurons exert feedback inhibition onto CA1 pyramidal cell dendrites (Klausberger and Somogyi, 2008). We recently showed that LTS neurons could be further divided into two distinct classes: Oriens Lacunosum Moleculare (OLM) neurons and Low Threshold High I_h_ (LTH) neurons (Hewitt et al., 2021). In this study, we used patch clamp electrophysiology to investigate differences in CA1 stratum oriens interneurons between wild type and FX mice. We found no significant differences in input resistance and action potential firing frequency in FS and OLM neurons between wild type and FX mice. In contrast, input resistance and action potential firing frequency were significantly lower in LTH neurons from FX mice compared to wild type. Previous studies showed that the intrinsic excitability of excitatory neurons is altered by changes in HCN channels (I_h_) in a cell-type specific manner in FX mice (Brager et al., 2012; Zhang et al., 2014; Kalmbach et al., 2015; Brandalise et al., 2020). Given that LTH neurons have a relatively higher expression of HCN channels (I_h_) compared to OLM cells (Hewitt et al., 2021), we tested the hypothesis that higher I_h_ contributes to the lower intrinsic excitability of FX LTH neurons. In agreement with this hypothesis, we found that the effect of I_h_ blocker ZD7288 on LTH neuron input resistance was higher in FX mice compared to wild type. Furthermore, direct measurement of I_h_ using voltage-clamp recordings found that the amplitude of I_h_ in LTH neurons was higher in FX compared to wild type.

## Methods

### Slice preparation

The University of Texas at Austin Institutional Animal Care and Use Committee approved all animal procedures. Mice had free access to food and water and were housed in a reverse light-dark cycle of 12 hr on/ 12 hr off. Experiments used male 2–4-month-old wild type and Fmr1 knockout (FX) mice on a C57/Bl6 background. Mice were anesthetized using a ketamine/xylazine cocktail (100/10 mg/kg) and then underwent cardiac perfusions with ice-cold saline consisting of (in mM): 2.5 KCl, 1.25 NaH_2_PO_4_, 25 NaHCO_3_, 0.5 CaCl_2_, 7 MgCl_2_, 7 dextrose, 205 sucrose, 1.3 ascorbate acid, and 3 sodium pyruvate (bubbled constantly with 95% O_2_/5% CO_2_ to maintain pH at ∼7.4). The brain was removed and sliced into 300 µM parasagittal sections from the middle hippocampus using a vibrating tissue slicer (Vibratome 300, Vibratome Inc). The slices were placed in a chamber filled with artificial cerebral spinal fluid (aCSF) consisting of (in mM): 125 NaCl, 2.5 KCl, 1.25 NaH_2_PO_4_, 25 NaHCO_3_, 2 CaCl_2_, 2 MgCl_2_, 10 dextrose, 1.3 ascorbic acid and 3 sodium pyruvate (bubbled constantly with 95% O_2_/5% CO_2_) for 30 minutes at 35°C and then held at room temperature until time of recording.

### Electrophysiology

Slices were placed in a submerged, heated (33-34 C°) recording chamber and continually perfused with aCSF (in mM): 125 NaCl, 3 KCl, 1.25 NaH_2_PO_4_, 25 NaHCO_3_, 2 CaCl_2_, 1 MgCl_2_, 10 dextrose, and 3 sodium pyruvate (bubbled constantly with 95% O_2_/5% CO_2_). Ionotropic glutamatergic and GABAergic synaptic transmission were blocked with 20 µM DNQX, 25 µM D-AP5, and 2 µM gabazine. Interneurons within the stratum oriens region of CA1 in slices were visualized with a Zeiss AxioScope or AxioExaminer under 60x magnification.

Current clamp recordings were made using a Dagan BVC-700 amplifier and custom written acquisition software using Igor Pro (WaveMetrics) or Axograph X (Axograph). Data were sampled at 20-50 kHz, filtered at 3 kHz, and then digitized by an InstruTECH ITC-18 interface (HEKA). The internal recording solution consisted of (in mM): 135 K-gluconate, 10 HEPES, 7 NaCl, 7 K_2_-phosphocreatine, 0.3 Na-GTP, 4 Mg-ATP (pH corrected to 7.3 with KOH). In some cases, 0.3% neurobiotin was added to the recording solution for post-hoc morphological reconstruction (see below). Recording electrodes were pulled from borosilicate glass and had an open tip resistance of 4-6 MΩ. Series resistance was compensated using the bridge balance circuit and was monitored throughout the experiment. Recordings in which series resistance exceeded 35 MΩ were excluded.

Prior to voltage clamp recordings, a train of action potentials was elicited using current clamp to first classify the cell as a FS, OLM, or LTH neuron. Following classification, slices were perfused with aCSF containing (in mM): 0.001 tetrodotoxin, 10 tetraethylammonium chloride, 5 4-aminopyridine, 0.1 BaCl_2_, 0.1 CdCl_2_, and 0.05 NiCl_2_ to block voltage-gated sodium, potassium, and calcium channels. Voltage clamp data were acquired using an Axopatch 200B amplifier (Molecular Devices) with custom written acquisition software in Igor Pro (WaveMetrics). Data were digitized at 20 kHz and filtered at 3 kHz. Inward currents were recorded in response to a series of 1000 ms hyperpolarizing voltage commands (-60 mV to -130 mV in -10 mV increments) from a holding potential of -30 mV.

### Drugs

All drugs were obtained from Tocris, Abcam pharmaceutical, or Sigma. Drugs were prepared from a 1000x stock solution in water.

### Neuronal Reconstruction and Analysis

Interneurons that were recorded with 0.3% neurobiotin (Vector Laboratories) in the pipette were processed with DAB for neuronal reconstructions. At the end of recording, the slices were fixed in 3% glutaraldehyde for at least 24hr. Slices underwent a servies of washes in 0.1 M phosphate buffer (PB) and then placed in 0.5%N H_2_O_2_ for 30 minutes. Slices were washed again in PB and then incubated in ABC (Vector Laboratories) solution for 24-48hr at 4°C. Slices were washed in PB before incubating in DAB solution (Vector Laboratories) for 1hr. H_2_O_2_ was slowly added to the slices every 10 minutes until a visible color change occurred. Slices were prepared for mounting by undergoing dehydration with ascending dilutions of glycerol (20%, 40%, 60%, 80%, 90%) for 10 minutes for each dilution until being placed in 100% glycerol solution. Slices were then mounted and identifieable interneurons were reconstructed with Neurolicida 6.0 software (MicroBrightField) at 40x maginification using a Leitz Diaplan microscope.

### Data analysis and statistics

Custom written software for either Igor Pro 7 or Axograph was used to analyze all physiology data. Input resistance was calculated from the linear portion of voltage-current relationship in response to a family of 1.5 s current injections of -40 pA to 40 pA in steps of 10 pA. Action potential firing frequency was calculated from the number of action potentials during a family of 1.5 s depolarizing current steps from 0 pA to 250 pA in steps of 25 pA. Action potential threshold was determined as the voltage where the first derivative exceeded 20 mV ms^-1^. The interspike interval (ISI) was defined as the distance in time between threshold of the first AP and the subsequent AP. The interspike interval ratio (ISI ratio) was calculated but dividing the time between the last two APs and the first two APs. All the statistical tests (unpaired t-tests, ANOVA, repeated measures of ANOVA, and Pearson’s R) were performed using Prism (GraphPad).

### Immunohistochemistry

Mice were deeply anesthetized using a ketamine/xylazine cocktail (100/10 mg/kg) and perfusion-fixed with ∼20 mL of cutting saline (see above) followed by ∼25 mL 4% paraformaldehyde in cutting saline at 4°C. The brain was removed, postfixed in PFA for 2 hours at RT, and placed in 30% sucrose in PBS overnight at 4°C. Horizontal thin sections (50 µm) containing the middle hippocampus were made on a cryostat (Leica) and placed in PBS. First, slices underwent antigen retrieval and were heated to between 85 °C and 95 °C for 30 minutes in 0.01M sodium citrate (pH adjusted to 6.0 with HCl) (Sigma). Slices were moved to a new plate and washed in 0.02% PBS-T 3 x 5 minutes. Slices were then incubated in a blocking buffer solution (5% normal goat serum, 0.2% triton+PBS) at RT for 1 hr, followed by primary antibodies: rabbit anti-parvalbumin (1:500, Swant) or rabbit anti-somatostatin (1:500, ImmunoStar) and mouse anti-FMRP (1:2 Developmental Studies Hybridoma Bank) in blocking solution for 48 hr at 4°C. Sections were then washed with 1x PBS 3 x 10 min and placed in blocking solution containing secondary antibodies (1:500 488 anti-mouse, 1:500 594 anti-rabbit) and Hoerst (1:2000) for 24 hr at 4°C. The slices were washed 4 x 10 min in 1x PBS and mounted on microscope slides and cover slipped with Fluormount-g (Invitrogen).

### Imaging and Image Analysis

The labelled sections were visualized on a Zeiss AxioImager fluorescent microscope with an ApoTome at 20x for analysis images and 40x for representative cell images. The images were captured by CCD camera, acquired by Stereo Investigator software (MBF Bioscience), and analyzed using Image J (NIH). Images were taken of the Stratum Oriens CA1 area of the hippocampus defined as the area between the CA1 pyramidal layer and the alveus and in between the subiculum and CA2/CA3. Co-expression analysis was completed in the SO CA1 using the cell counter software in Image J (NIH). Neurons were first counted and traced in the 594 channel to identify PV+ and SOM+ neurons, and then overlayed in the 488 channel to identify co-expression of FMRP. Cells were counted if they co-expressed the interneuron marker (PV or SOM) and FMRP in stratum oriens of area CA1. Slices from 4 wild type mice across 15 sections were included in the analysis. In total, there were 171 PV and 174 SOM positive neurons that were counted for analysis, we then took the percentage of PV and SOM positive neurons that also expressed FMRP.

## Results section

### LTS neuron intrinsic excitability is lower in FX mice

We made whole-cell current clamp recordings from neurons in the stratum oriens of area CA1 in the hippocampus from wild type (WT) and Fmr1 knock-out (FX) mice. After noting the resting membrane potential, neurons were held at -70 mV and two families of current injections were used to identify the type of interneuron. First, a series of 1-sec long depolarizing current injections was used to measure action potential firing. Second, a series of small subthreshold current injections to determine the input resistance (R_N_).Fast spiking (FS) interneurons were identified by having a maximum action potential firing rate >100 Hz, no spike frequency adaptation, and low R_N_. Low threshold spiking (LTS) interneurons by contrast had a maximum action potential firing rate <80Hzreadily apparent spike frequency adaptation, and high R_N_.

Voltage responses to hyperpolarizing current injections in FS interneurons from both wild type and FX mice exhibited no sag or rebound and had comparatively low input resistances (<180 MΩ; Figure 1A). We found no differences in the RMP (unpaired t-test; df = 15; t=1.834; p=0.08) or R_N_ (unpaired t-test; df = 14; t=0.065; p=0.52) between wild type and FX FS interneurons (Figure 1B-C). FS neurons exhibited the expected high frequency firing rate and no adaptation in wild type and FX mice (Figure 1D). A plot of action potentials fired as a function of current amplitude revealed no difference in the number of action potentials elicited across a range of depolarizing current amplitudes between wild type and FX FS interneurons (two-way ANOVA, main effect of genotype F(1,29)=1.107; p=0.301; Figure 1E).

**Figure 1:**
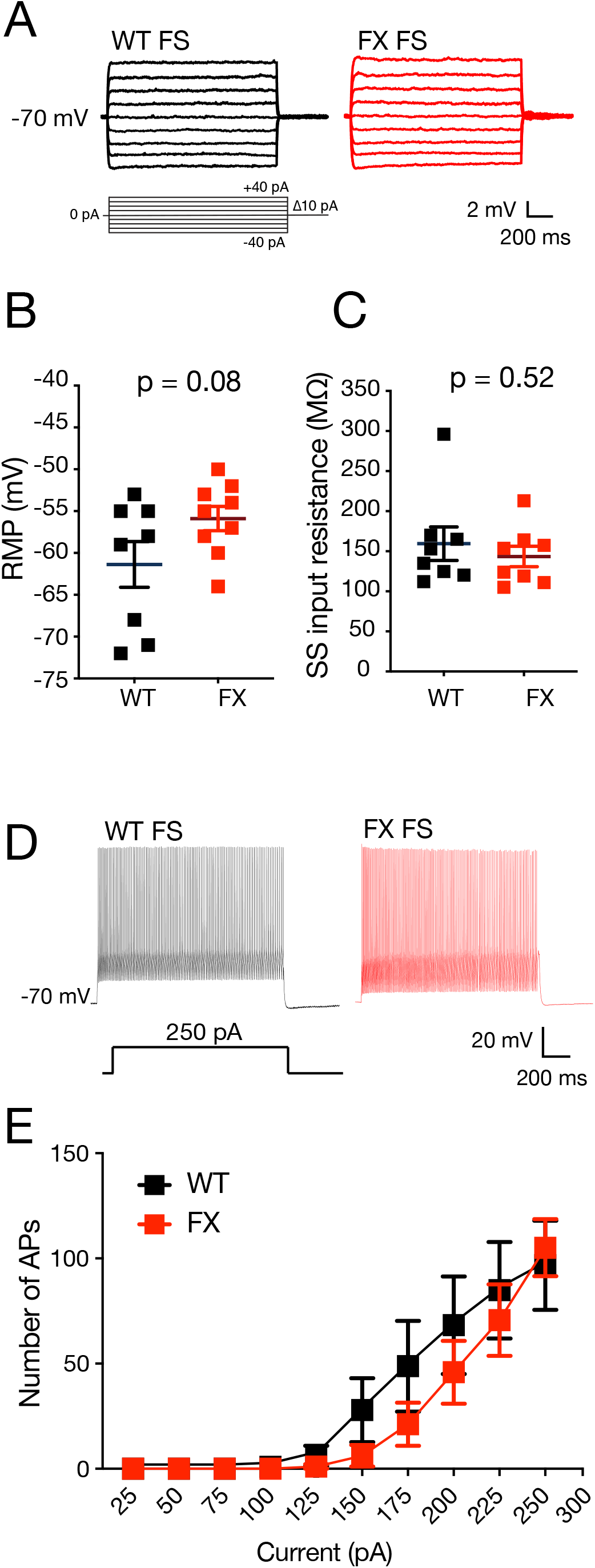
Subthrehsold properties and action potential firing for fast spiking interneurons in stratum oriens between WT and FX mice. a) Voltage responses to a family of small current steps in WT and FX FS neurons. b) Resting membrane potential is not different between WT and FX FS interneurons (unpaired t-test; df = 15; t=1.834; p=0.08) c) Steady state input resistance is not different between WT and FX FS interneurons (unpaired t-test; df = 14; t=0.065; p=0.52) d) Action potential firing in WT and FX FS interneurons in response to a 250 pA current injection. e) There was no difference in action potential firing output across a range of depolarizing current steps between WT and FX neurons (two-way ANOVA, main effect of genotype F(1,29)=1.107; p=0.301).

In contrast to FS interneurons, subthreshold voltage responses from LTS interneurons exhibited a prominent sag and rebound in response to current injections (Figure 2A), and a comparatively high input resistance (range: 174-740 MΩ). Although there was no difference in RMP between wild type and FX LTS neurons (unpaired t-test; df = 54; t=0.69; p=0.49; Figure 2B), FX LTS neurons had a significantly lower R_N_ compared to wild type (unpaired t-test; df = 53; t=2.622; p=0.01; Figure 2C). As expected, both wild type and FX hippocampal LTS neurons displayed spike frequency adaptation (Figure 2D). When compared across a range of depolarizing current steps, FX LTS neurons fired significantly fewer action potentials compared to wild type (two-way ANOVA, main effect of genotype F(1,50)=5.69; p=0.02; Figure 2E). These data therefore suggest that the intrinsic excitability of LTS, but not FS, interneurons is lower in FX mice.

**Figure 2:**
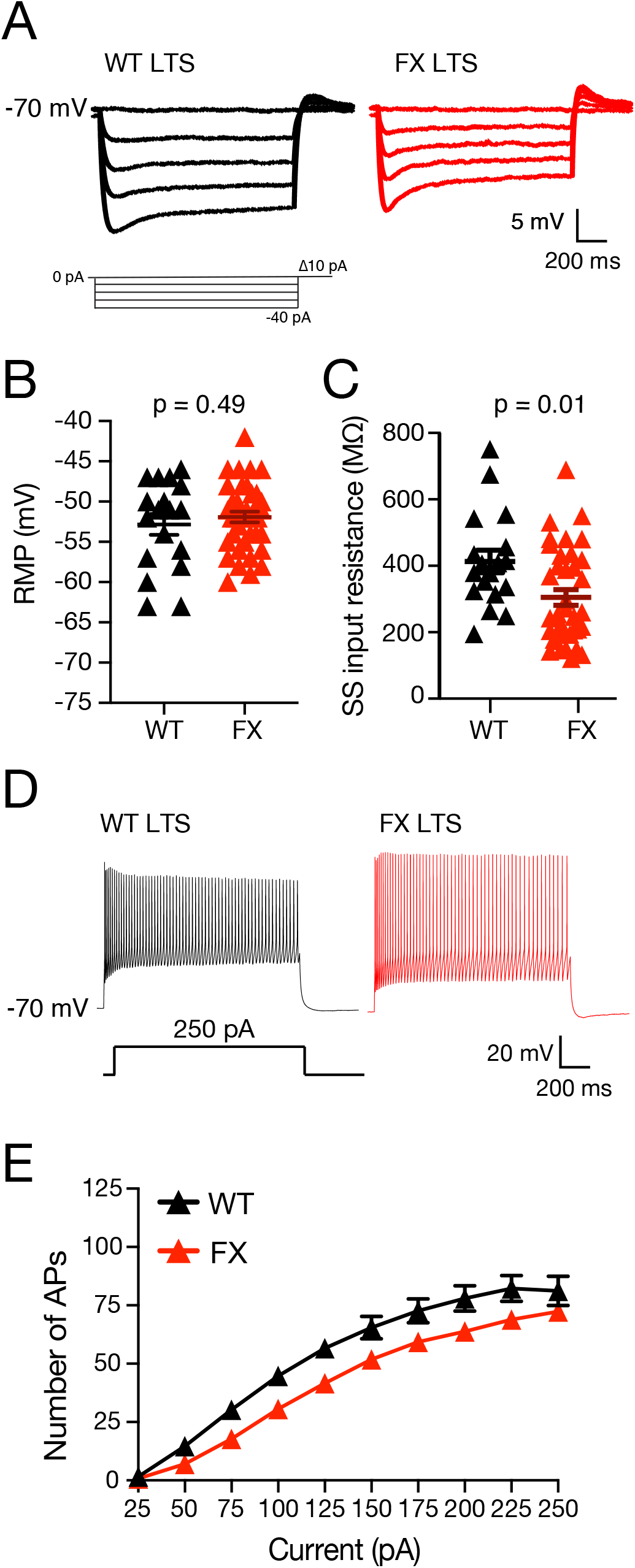
Input resistance and action potential firing are lower in low-threshold spiking interneurons in stratum oriens of FX mice compared to WT. a) Voltage responses to a family of hyperpolarizing current steps in WT and FX LTS interneurons. b) Resting membrane potential is not different between WT and FX LTS interneurons (unpaired t-test; df = 54; t=0.69; p=0.49). c) Input resistance is lower in FX LTS interneurons compared to WT (unpaired t-test; df = 53; t=2.622; p=0.01) d) Action potential firing in WT and FX LTS interneurons in response to a 250 pA current injection. e) FX LTS neurons fire fewer action potentials compared to WT across a range of depolarizing current steps (repeated measures ANOVA, main effect of genotype (F (1,50) = 5.69; p = 0.02)).

### Both FS and LTS interneurons co-express FMRP

Given that we observed differences in both R_N_ and action potential firing in LTS interneurons, but not FS interneurons, between wild type and FX mice, we wanted to test if there were differences in FMRP expression between FS and LTS interneurons. One of the canonical methods of identifying interneuron subtypes is the expression of specific proteins. Somatostatin (SOM) and parvalbumin (PV) are established markers of hippocampal LTS and FS interneuron subtypes respectively ((Baude et al., 2007; Tricoire et al., 2011; Chittajallu et al., 2013) but see (Hu et al., 2013; Pelkey et al., 2017)). We used immunohistochemistry to identify the co-expression of FMRP with either SOM-positive or PV-positive neurons in stratum oriens of area CA1 in wild type mice (Figure 3A-B). We found that the percentage of SOM+ (putative LTS) neurons which co-express FMRP is greater compared to PV+ (putative FS) neurons) (unpaired t-test; df = 6; t=5.119; p=0.002; Figure 3C). Our analysis indicates SOM interneurons more reliably express FMRP compared to PV interneurons.

**Figure 3:**
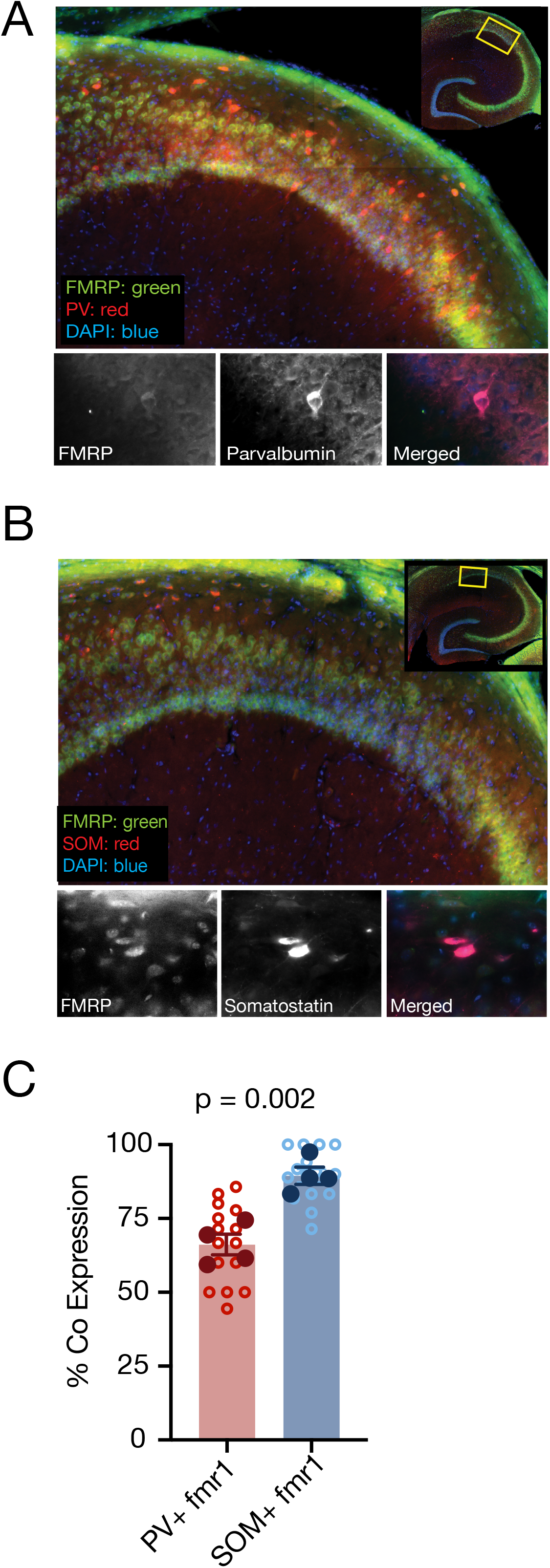
Co-expression of FMRP with parvalbumin and somatostatin in stratum oriens of the hippocampus. a) Immunohistochemical labeling of FMRP (green) and parvalbumin (red) in the CA1 region of the hippocampus. DAPI stain is shown in blue. Inset, low magnification image with box indicating the analyzed region. Individual PV+ cell shown below at 40x magnification with FMRP expression. b) Immunohistochemical labeling of FMRP (green) and somatostatin (red) in the CA1 region of the hippocampus. DAPI stain is shown in blue. Inset, low magnification image with box indicating the analyzed region. Individual SOM+ cell shown at 40x magnification with FMRP expression. c) Percent of FMRP+ neurons that co-express either PV or SOM (interneuron marker / FMRP+ neurons). In stratum oriens of CA1, SOM+ neurons show more co-expression with FMRP compared to PV+ neurons (PV images n= 4 mice, 14 slices, SOM images n= 4 mice, 14 slices unpaired t-test; df = 6; t=5.119; p=0.002).

### Both subclasses of LTS neurons are present in FX mice

We recently demonstrated that LTS interneurons in stratum oriens of area of CA1 can be separated into at least two distinct groups, oriens lacunosum-moleculare cells (OLM) and low threshold high I_h_ cells (LTH) (Hewitt et al., 2021). One reliable measure for separating OLM and LTH cells is the degree of spike frequency adaptation (SFA). LTH neurons have more pronounced SFA compared to OLM neurons (Hewitt et al., 2021). Using our previously published criteria (Hewitt et al., 2021), we separated LTS interneurons into OLM and LTH groups (Figure 4A). Consistent with our prior criteria, we separated LTS cells into LTH and OLM neurons based on the ISI ratio (Figure 4B). Spike frequency adaptation (see Methods) was not different between wild type and FX mice for both LTH cells (unpaired t-test; df = 22; t=0.536; p=0.51; Figure 4C) and OLM cells (unpaired t-test; df = 30; t=0.662; p=0.59; Figure 4D). We found OLM and LTH neurons in roughly the same proportions between wild type and FX mice (Figure 4E).

**Figure 4:**
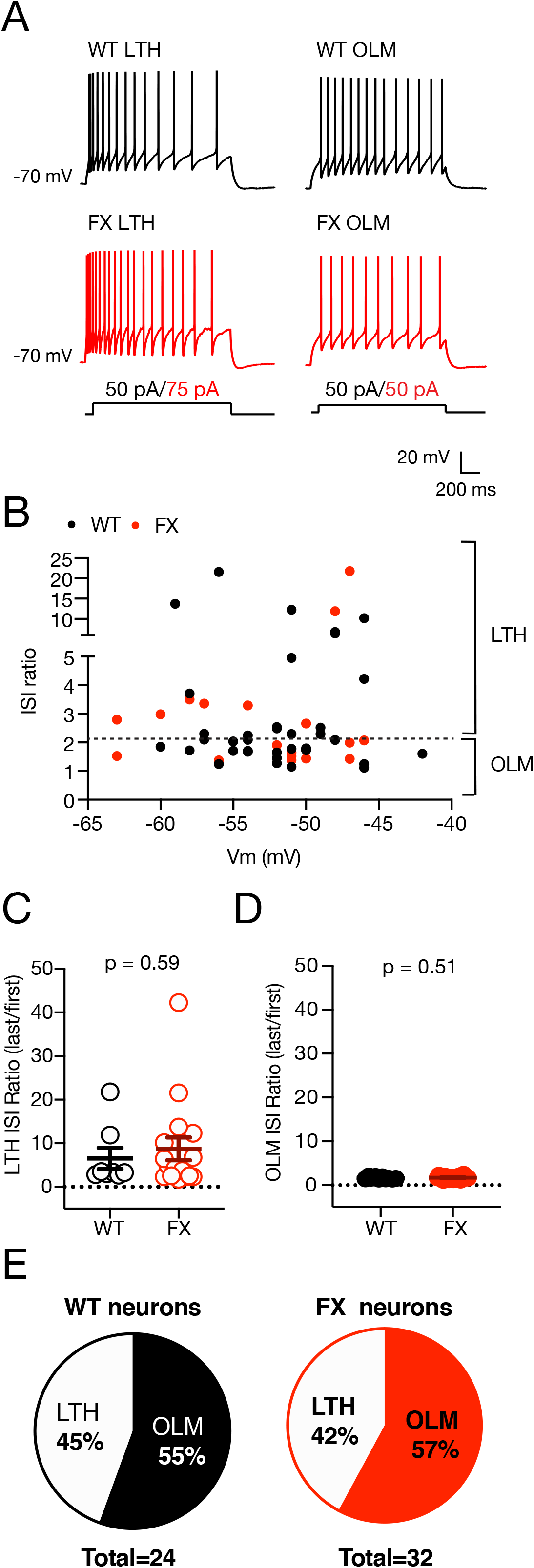
Both OLM and LTH cell types of low-threhsold spiking interneurons are found in the FX hippocampus. a) Voltage traces from the first depolarizing current step to elicit action potentials in LTH and OLM neurons from WT (black) and FX (red) mice. b) ISI ratio does not have a significant relationship to resting membrane potential for WT and FX LTS interneurons. Dashed line indicates the ISI ratio cut off between LTH and OLM neurons (Hewitt et al., 2021). c-d) ISI ratio is not different between WT and FX mice for LTH neurons (d, unpaired t-test; df = 22; t=0.536; p=0.59) or OLM (e, unpaired t-test; df = 30; t=0.662; p=0.51). e) The proportion of OLM and LTH neurons is not different between WT and FX mice (total = the number recorded cells).

### LTH interneurons have lower intrinsic excitability in FX mice

LTH interneurons have a lower R_N_ and more prominent sag and rebound compared to OLM interneurons consistent with higher I_h_ and our previous work (Hewitt et al., 2021), (Figure 5A and Table 1). FX LTH neurons had a lower R_N_ compared to wild type LTH neurons (unpaired t-test; df = 22; t=3.575; p=0.001; Figure 5B). Surprisingly, there was no difference in RMP, sag, or rebound between wild type and FX LTH neurons (Table 1). We found no differences in RMP, R_N_, sag, or rebound between wild type and FX OLM interneurons (Figure 5C-D, Table 1). In response to depolarizing current injections, FX LTH interneurons fired significantly fewer action potentials compared to wild type (two-way ANOVA, main effect of genotype F(1,19)=7.971; p=0.01; Figure 6A-B). In contrast, there was no significant difference in the number of action potentials fired between wild type and FX OLM interneurons (two-way ANOVA, main effect of genotype F(1,30)=0.83; p=0.369; Figure 6C-D).

**Figure 5:**
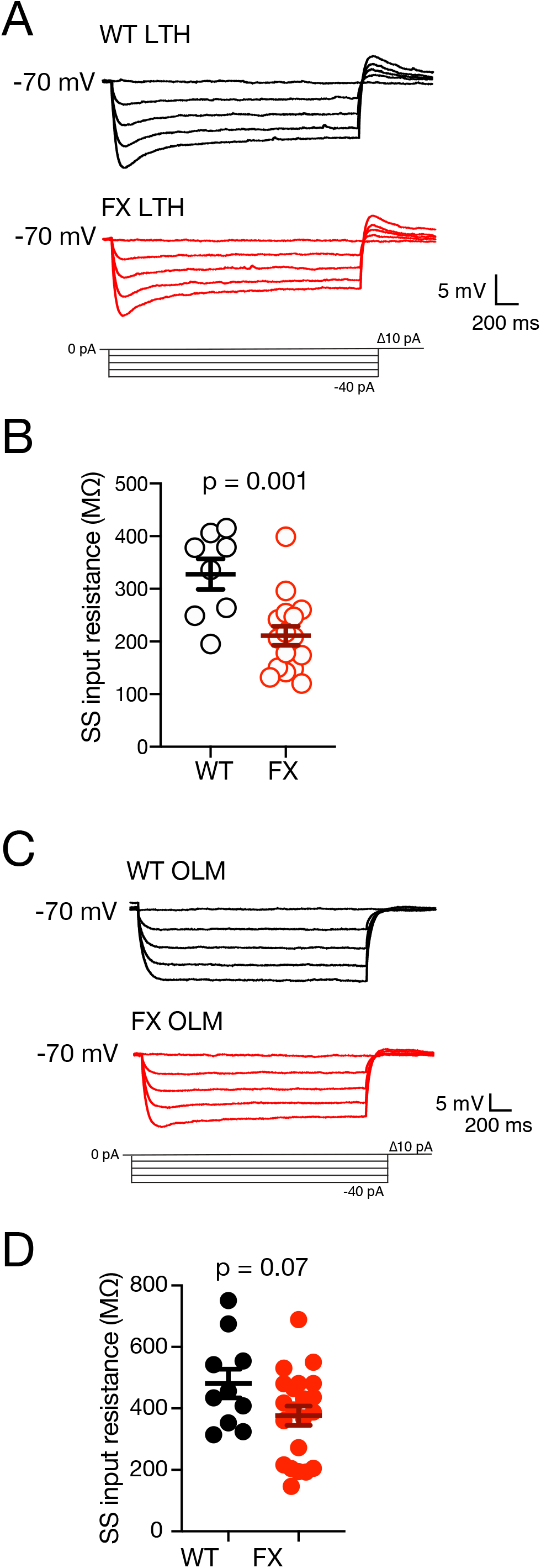
LTH, but not OLM, neurons have a lower input resistance in FX mice compared to WT. a) Voltage traces of CA1 LTH WT and FX LTH neurons in response to family of hyperpolarizing current steps. b) FX LTH neurons show a lower steady state input resistance compared to WT LTH neurons (unpaired t-test; df = 22; t=3.575; p=0.001). c) Voltage traces of WT and FX OLM neurons in response to a family of hyperpolarizing current steps. d) no differences in steady-state input resistance of WT and FX OLM cells (unpaired- t-test, df =29, t=1.87, p= 0.07).

**Figure 6:**
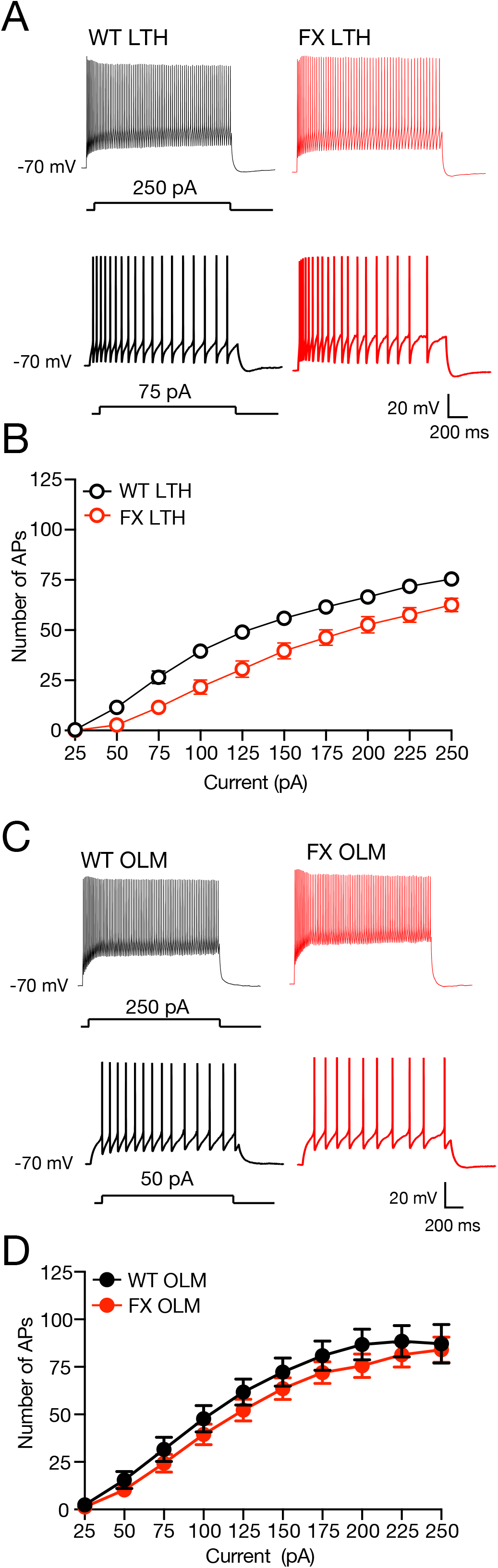
LTH, but not OLM, neurons fire significantly fewer APs in FX mice compared to WT. a) Representative voltage responses to a 75 pA (top) and 250 pA (bottom) current step in WT and FX LTH neurons. b) FX LTH neurons fired significantly fewer action potentials across a range of current injections compared to WT (repeated measures ANOVA, main effect of genotype (F(1,19) = 7.971; p = 0.01). c) Representative voltage responses to a 50 pA (top) and 250 pA (bottom) current injection in WT and FX OLM interneurons. d) There was no significant difference in action potential firing output between WT and FX OLM interneurons (two-way ANOVA, main effect of genotype F(1,30)=0.83, p=0.369).

**Table 1:** No difference in sag and rebound response between wild type and FX LTH or OLM neurons. Resting membrane potential (RMP), sag, and rebound responses of wild type and FX LTH and OLM neurons. No difference in RMP, sag or rebound response between wild type and FX LTH neurons (RMP: unpaired- t-test, df =22, t=1.49, p= 0.14; sag: unpaired t-test, df=18, t=0.44, p=0.65; rebound: unpaired t-test, df=16, t=0.42, p=0.67) No difference in sag or rebound response between wild type and FX OLM neurons (RMP: unpaired- t-test, df =30, t=0.45, p= 0.65; sag: unpaired t-test, df = 29, t=1.52, p=0.14; rebound: unpaired t-test, df = 11, t=2.13, p=0.06).

|  | wild type | FX | p-value |
| --- | --- | --- | --- |
| **LTH**<br>(n = 8 wt, 16 ko) |  |  |  |
| RMP | -54 ± 2.04 | -51 ± 2.04 | 0.14 |
| Sag | 1.23 ± 0.05 | 1.20 ± 0.05 | 0.65 |
| Rebound | -0.28 ± 0.06 | 0.31 ± 0.06 | 0.67 |
| **OLM**<br>(n = 10 wt, 22 ko) |  |  |  |
| RMP | -51 ± 1.71 | -52 ± 1.71 | 0.65 |
| Sag | 1.04 ± 0.05 | 1.12 ± 0.05 | 0.14 |
| Rebound | -0.04 ± 0.01 | -0.07 ± 0.01 | 0.06 |

### Action potential threshold is not different in LTH neurons

We hypothesize that the lower input resistance in FX LTH neurons accounts for the reduced action potential firing rate. However, changes in action potential threshold were reported in multiple cell types in FX mice (Deng and Klyachko, 2016; Kalmbach and Brager, 2020; Deng et al., 2021). To determine if action potential threshold was different, we analyzed the first elicited action potential between wild type and LTH neurons. There was no difference in threshold for the first action potential in LTH neurons between wild type and FX mice (Figure 7A-B; unpaired- t-test, df =21, t=1.02, p= 0.31). Further analysis of the first action potential did not find any differences in amplitude, half-width, afterhyperpolarization, maximum rate of rise, or maximum rate of decay (Table 2). During repetitive firing, the threshold for action potential firing becomes more depolarized due changes in the availability for voltage-gated sodium channels (e.g., cumulative inactivation) (Colbert and Pan, 2002). Thus, it is possible that although the threshold for the first action potential is not different, that the change in action potential threshold during repetitive firing is different between wild type and FX LTH neurons. Consistent with previous results, we found that action potential threshold depolarized during a train of 25 action potentials. There was, however, no difference in the change in action potential threshold during repetitive firing between wild type and FX mice (Figure 7C; two-way ANOVA, main effect of genotype F(1,16)=0.53; p=0.47). We found the same lack of difference in action potential threshold for OLM neurons (Figure 7D-E; (unpaired t-test, df=18 t=0.28 p= 0.78); (two-way ANOVA, main effect of genotype F(1,13)=0.02, p=0.88).

**Figure 7:**
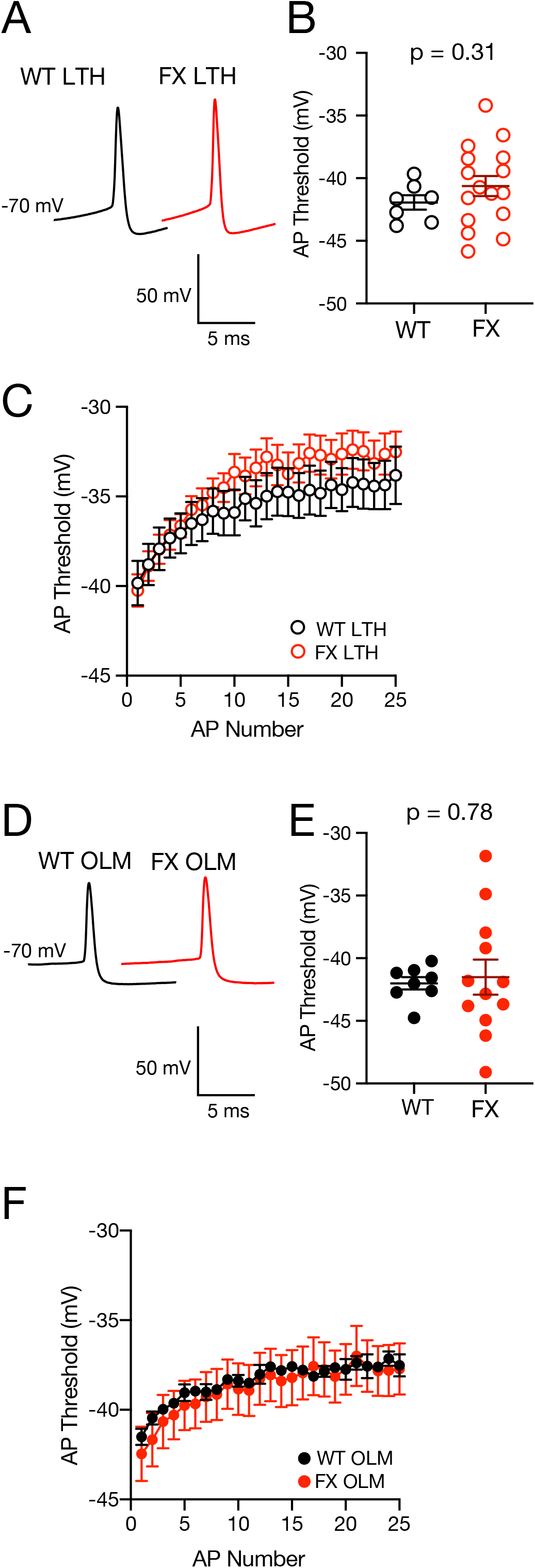
Action potential threshold in LTH and OLM neurons is not different between WT and FX mice. a) Representative voltage traces of the first action potential elicited by the smallest current injection in WT and FX LTH neurons. b) AP threshold is not different between WT and FX LTH neurons (unpaired- t-test, df =21, t=1.02, p= 0.31). c) The change in action potential threshold during a train of 25 action potentials is not different between WT and FX LTH neurons (two-way ANOVA, main effect of genotype F(1,16)=0.53; p=0.47). d) Representative voltage traces of the first action potential elicited by the smallest current injection in WT and FX OLM neurons. e) AP threshold is not different between WT and FX OLM cells (unpaired t-test, df=18 t=0.28 p= 0.78) f) The change in action potential threshold during a train of 25 action potentials is not different between WT and FX OLM neurons (two-way ANOVA, main effect of genotype F(1,13)=0.02, p=0.88).

**Table 2:** Action potential properties in LTH neurons are not different between wild type and FX mice. Analysis of single AP properties showed no difference between wild type and FX LTH neurons. AP Amplitude: (unpaired t-test, df = 16, t=1.17, p=0.26); AP half-width: (unpaired t-test, df = 16, t=0.71, p=0.49); AHP: (unpaired t-test, df = 16, t=0.15, p=0.88; Max rate of rise: (unpaired t-test, df = 16, t=0.75, p=0.46); Max rate of fall: (unpaired t-test, df = 16, t=0.52, p=0.58).

|  | wild type (n=11) | FX (n=7) | p-value |
| --- | --- | --- | --- |
| AP amplitude (mV) | 101.7 ± 3.03 | 98.1 ± 3.03 | 0.26 |
| AP half-width (ms) | 0.76 ± 0.06 | 0.71 ± 0.06 | 0.49 |
| AHP (mV) | -21.1 ± 2.6 | -20.89 ± 2.6 | 0.88 |
| Max. rate of rise<br>(mV/ms) | 285.6 ± 21.4 | 269.5 ± 21.4 | 0.46 |
| Max. rate of fall<br>(mV/ms) | -137 ± 13.4 | -144.4 ± 13.4 | 0.58 |

### Neuronal morphology is not different between wild type and FX LTH neurons

CA1 LTH neurons in wild type mice have oblong cell bodies located in stratum oriens with dendrites extending along the CA3 subicular axis (Hewitt et al. 2021). We reconstructed 4 wild type (Figure 8A) and 3 FX LTH (Figure 8B) neurons to determine changes in gross neuronal morphology. Wild type and FX LTH neurons had similar gross neuronal morphology. We found no differences in dendrite number, somato-dendritic length, or cell surface area between wild type and FX LTH neurons (Figure 8C-E).

**Figure 8:**
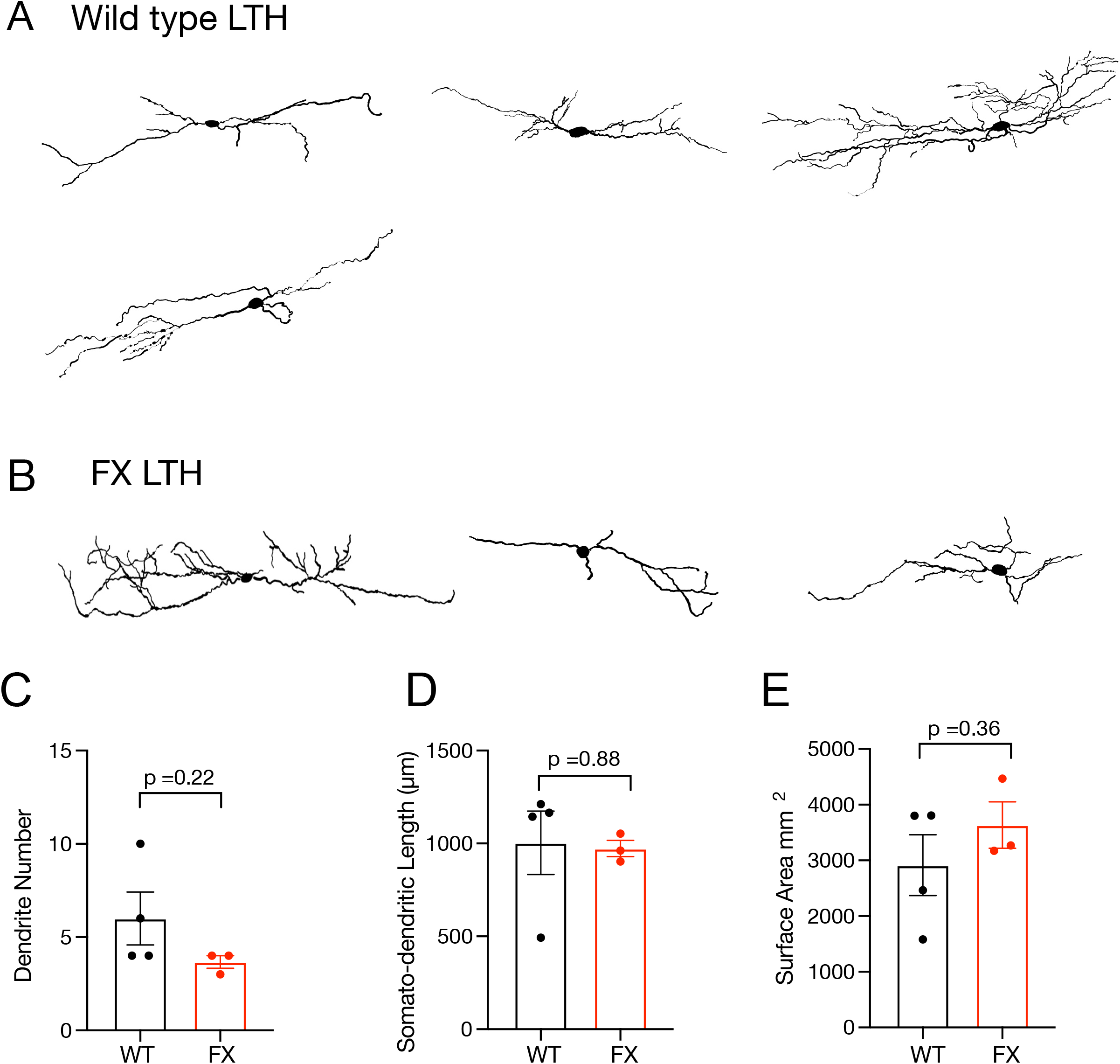
Neuronal morphology is not different between WT and FX LTH neurons. a,b) Neurolicida morphological reconstructions of WT (a) (n=4 mice; 4 cells) and FX LTH (b) neurons (n=3 mice; 3 cells). c-e) There was no difference in the number of dendritic branches (c, unpaired t-test, df=5 t=1.37 p= 0.22), somato-dendritic length (d, unpaired t-test, df=5 t=0.15 p= 0.88), or surface area (e, unpaired t-test, df=5 t=0.98 p= 0.36) between WT and FX LTH neurons.

### Block of I_h_ normalizes input resistance between wild type and FX LTH neurons

Both excitatory and inhibitory neurons express HCN channels (I_h_) which contribute to the resting properties of the neuron (Maccaferri and McBain, 1996; Magee, 1999; Santoro et al., 2000; Lupica et al., 2001; Poolos et al., 2002). We previously showed that LTH neurons have a higher contribution of I_h_ to the R_N_ compared to OLM cells (Hewitt et al., 2021). To test the hypothesis that higher I_h_ contributes to the lower R_N_ in FX LTH neurons we made current clamp recordings from wild type and FX LTH interneurons before and after extracellular application of 50 µM ZD7288 to block HCN channels (Figure 9A). Consistent with our results in Figure 6, the baseline R_N_ of FX LTH interneurons was significantly lower compared to wild type (unpaired t-test: t = 2.305, df = 11, p = 0.0416; Figure 9B). Given the significant contribution of I_h_ to R_N_ in LTH neurons, the block of I_h_ with ZD7288 significantly increased the R_N_ of LTH neurons in both wild type (paired t-test: t = 4.168, df = 6, p = 0.0059) and FX mice (paired t-test: t = 3.356, df = 5, p = 0.0202) (Figure 9A-B). Interestingly, R_N_ was not significantly different between wild type and FX LTH neurons after application of ZD7288 (unpaired t-test: t = 0.3221, df = 11, p = 0.7534; Figure 9B). These results support our hypothesis that FX LTH interneurons have more I_h_ compared to wild type.

**Figure 9:**
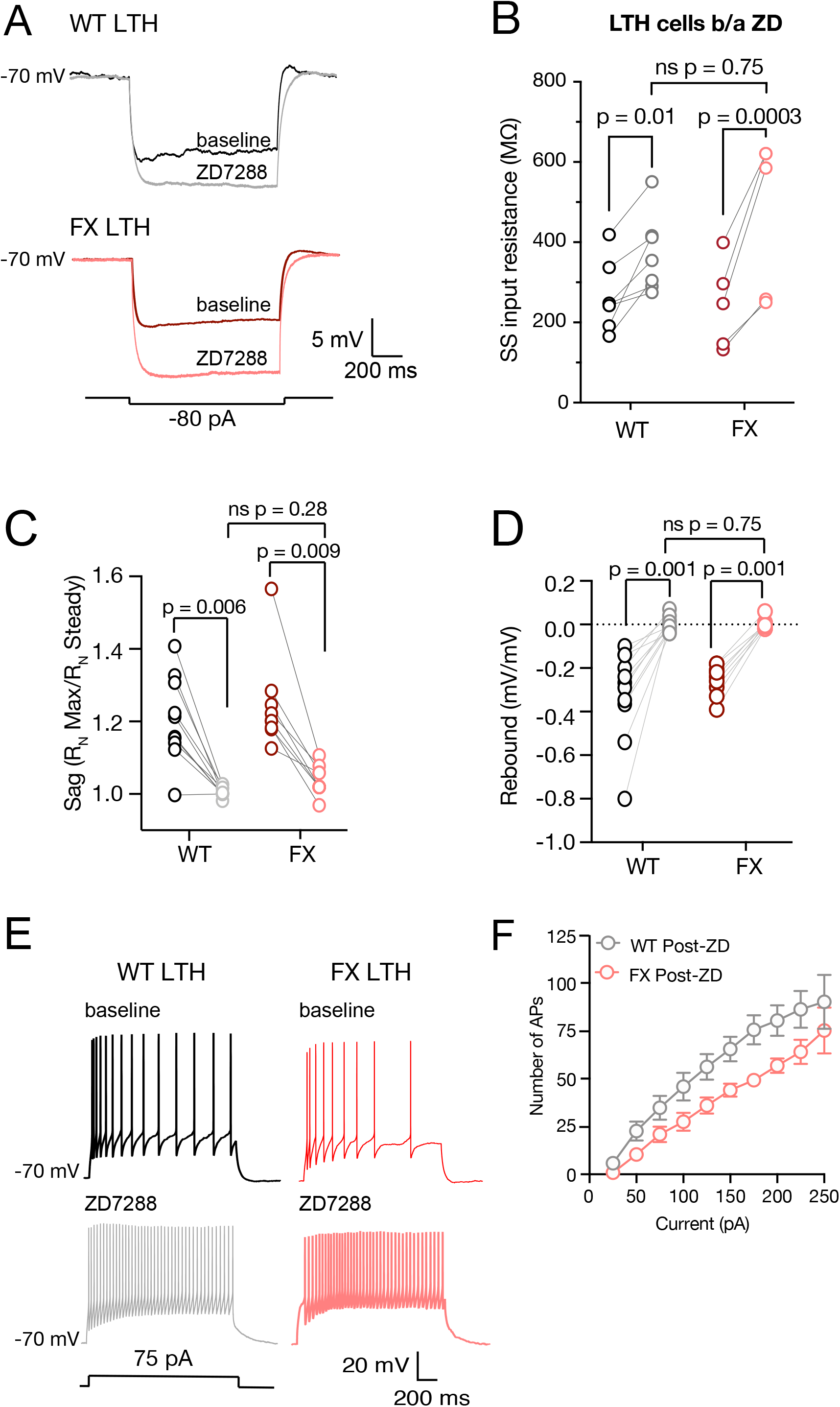
Block of I_h_ with ZD7288 normalizes input resistance but not AP firing between WT and FX LTH neurons. a) Representative voltage responses to a hyperpolarizing step of -80 pA before and after 50 µM ZD7288 for LTH neurons from WT (black, baseline; grey ZD7288) and FX (red, baseline; pink, ZD7288) mice. b) ZD7288 significantly increased steady-state input resistance in both WT and FX LTH cells (WT: unpaired *t*-test, p = 0.01; FX: unpaired *t*-test, p = 0.0003). Note that there is no significant difference in steady-state input resistance between WT and FX LTH neurons in input resistance after ZD7288 (unpaired *t-*test, p = 0.75). c) ZD7288 significantly decreases voltage sag in WT and FX LTH cells (WT: unpaired t-test, df=20 t=3.03 p= 0.006; FX: unpaired t-test, df=14 t=4.17 p= 0.009). Note that there was no difference in sag between WT and FX LTH neuron after ZD7288 (unpaired t-test, df=17 t=1.11 p= 0.28). d) ZD7288 significantly decreases rebound in both WT and FX LTH cells (WT: unpaired t-test, df=20 t=4.95 p= 0.001; FX: unpaired t-test, df=15 t=9.76 p= 0.001). Note that there was no difference in rebound between WT and FX LTH neuron rebound after ZD (unpaired t-test, df=17 t=0.32 p= 0.75). e) Representative actin potentials in responses to a depolarizing step of 75 pA before and after ZD7288 for LTH neurons from WT (black, baseline; grey ZD7288) and FX (red, baseline; pink, ZD7288) mice. f) FX LTH neurons significantly fewer action potentials than WT neurons after ZD application (two-way ANOVA, main effect of genotype F(1,17)=5.43, p=0.03).

Voltage sag and rebound response are current clamp measurements that can be indicative of I_h_ and can be significantly decreased with bath application of ZD7288. As expected, ZD7288 significantly decreased both voltage sag (WT: unpaired t-test, df=20 t=3.03 p= 0.006; FX: unpaired t-test, df=14 t=4.17 p= 0.009, Figure 9C) and rebound (WT: unpaired t-test, df=20 t=4.95 p= 0.001; FX: unpaired t-test, df=15 t=9.76 p= 0.001, Figure 9D) in wild type and FX LTH neurons.

We hypothesized that the lower input resistance underlies the lower action potential firing in FX LTH neurons. As ZD7288 normalized the input resistance between wild type and FX LTH neurons, we asked whether blocking I_h_ would similarly rescue action potential firing in FX LTH neurons. Surprisingly, FX LTH neurons post-ZD application fired significantly fewer APs comparted to wild type LTH neurons (Figure 9E-F, two-way ANOVA, main effect of genotype F(1,17)=5.43, p=0.03).

### LTH neurons in FX mice have higher I_h_

Given the effect of ZD7288 on input resistance, we used a previously published current clamp, voltage clamp protocol to directly measure I_h_ in LTH neurons (Maccaferri and McBain, 1996; Hewitt et al., 2021). First, we measured the ISI ratio of action potential firing using current clamp to identify the cell as FS, OLM, or LTH. Following cell classification, we applied a combination of pharmacological agents to block voltage-gated Na^+^, K^+^, and Ca^2+^ channels (Figure 10A). After 20 minutes, we switched to voltage clamp and measured the current in response to a family of hyperpolarizing voltage steps from a holding potential of -30 mV before and after application of 50 µM ZD7288. The currents in the presence of ZD was subtracted from the pre-ZD currents (Figure 10B-D). Consistent with the prominent sag and rebound, a slowly activating ZD-sensitive inward current was recorded from both wild type and FX LTH interneurons. We found that the ZD-sensitive current was larger in FX LTH neurons compared to wild type (two-way ANOVA, main effect of genotype: F(1,29)=1.107, p=0.0005; Figure 10D-E). These results suggest that the lower input resistance in FX LTH neurons is due in part to higher I_h_.

**Figure 10:**
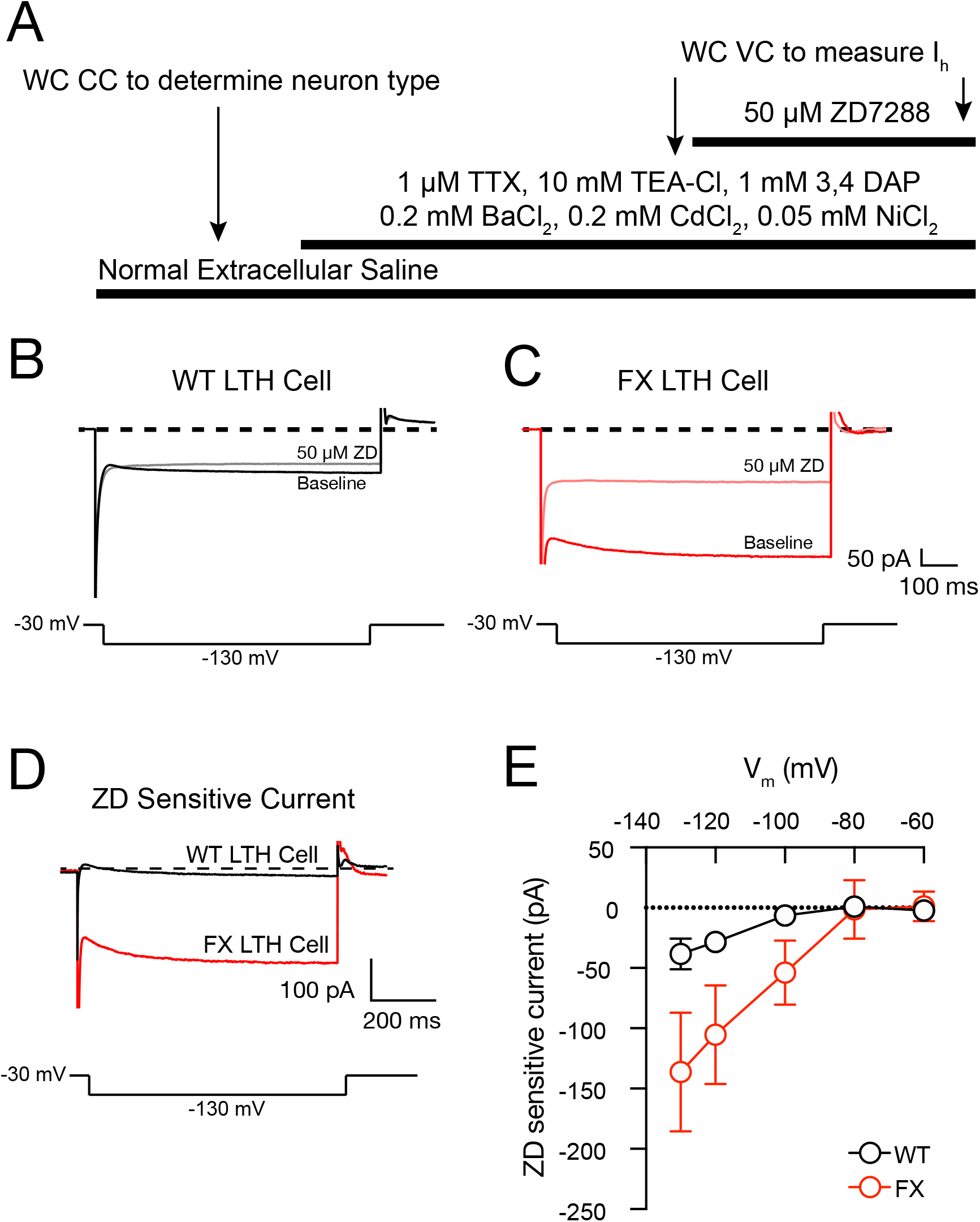
FX LTH neurons have more hyperpolarization activated current Ih compared to WT LTH neurons. a) Experimental procedure for measuring I_h_ in WT and FX LTH neurons. b) Current measured in response to a voltage step to -130 mV before and after 50 µM ZD7288 in WT LTH neuron (black, baseline; grey, ZD7288) and post-bath application of I_h_ channel blocker ZD7288 (grey). c) Current measured in response to a voltage step to -130 mV before and after 50 µM ZD7288 in FX LTH neuron (red, baseline; pink, ZD7288). d) representative traces of ZD sensitive current for a -130 mV step from WT (black) and FX (red) LTH neurons. e) FX LTH neurons have more I_h_ compared to WT (two-way ANOVA, main effect of genotype: F(1,29)=1.107, p=0.0005).

## Discussion

There are many studies investigating excitatory pyramidal neurons in FXS, while investigations of inhibitory interneurons are lacking. We investigated the intrinsic physiological properties of FS and LTS inhibitory interneurons in the stratum oriens of area CA1 in the hippocampus from wild type and FX mice. We found no differences in either subthreshold properties or action potential firing between wild type and FX FS interneurons. By contrast, FX LTS interneurons had a lower input resistance and fired fewer action potentials compared to wild type. We used the interneuron markers parvalbumin (PV) to label putative fast spiking interneurons and somatostatin (SOM) to label putative low threshold spiking interneurons and determine the co-expression of FMRP. Although both PV+ and SOM+ cells expressed FMRP, SOM+ neurons more reliably expressed FMRP compared to PV+ cells.

As described in our recent paper (Hewitt et al., 2021), we subdivided low threshold spiking interneurons into OLM and LTH interneurons. Current clamp experiments revealed FX LTH, but not OLM, interneurons had a significantly lower input resistance and fired fewer action potentials across a family of current steps. Current clamp experiments revealed that FX LTH neurons were more sensitive to the h-channel blocker ZD7288 suggesting that the differences in input resistance were due in part to higher I_h_. Voltage clamp experiments confirmed that FX LTH neurons have higher I_h_ compared to wild type.

Changes in voltage-gated ion channel function and expression, including HCN channels, were identified in FX excitatory pyramidal cells (Brager et al., 2012; Deng et al., 2013, 2019; Zhang et al., 2014; Kalmbach et al., 2015; Deng and Klyachko, 2016; Routh et al., 2017). Here we show FX LTH interneurons have a lower input resistance due in part to a larger h-channel current. While other stereotypical effects of higher I_h_ were not observed in FX LTH neurons (e.g., depolarized RMP, higher sag and rebound), this could be due in part to differences in the h-channel kinetics of h-channels and/or the localization of the channels (dendritic vs somatic expression) in FX LTH neurons. To properly investigate this question, cell-attached or outside-out patch clamp recordings would be necessary to address the space clamp limitations of whole-cell voltage clamp. Either of these techniques would be somewhat limited or difficult to interpret. First, given the requirement of cell type identification, cell-attached patch clamp would be problematic. Second, although outside-patch clamp is potentially feasible, visualization of and electrode placement on the small dendrites of stratum oriens interneurons would make this approach difficult.

Block of I_h_ with bath application of ZD7288 rescued the lower input resistance in FX LTH neurons. Despite this, LTH neurons in FX mice still fired significantly fewer action potentials compared to wild type in the presence of ZD7288. This result suggests that higher I_h_ is not solely responsible for altered intrinsic excitability in FX LTH neurons. Sodium and potassium channels are critical to generating the action potential waveform. Although changes in both Na+ and K+ channels were reported in both excitatory and inhibitory neurons in FX mice (Brown et al., 2010; Strumbos et al., 2010; Routh et al., 2013; Kalmbach et al., 2015; Deng and Klyachko, 2016) we did not observe any changes in the action potential waveform (Table 2) that would support significant changes in either Na^+^ or K^+^ channel function. Na^+^ and K^+^ channels also contribute spike frequency adaptation in neurons during repetitive firing by modulating both action potential threshold and interspike interval (Bean, 2007). We found no difference in action potential threshold or interspike interval ratio between wild type and FX LTH neurons during repetitive action potential firing. It is possible that differences in K^+^ channel function and/or expression contribute to the lower action potential firing in FX LTH neurons. Due to the incredible diversity of voltage-gated potassium channels and the very limited knowledge of interneuron specific potassium channels however, it would be exceeding difficult to isolate which channels may contribute to the lower AP number in FX LTH neurons (for review: (Lien et al., 2002; Pelkey et al., 2017)

We found no difference in the intrinsic excitability of hippocampal FS interneurons between wild type and FX mice. There are reports however, that found changes in FS interneurons located in other brain areas in FX mice. FS neurons in layer 4 of the somatosensory cortex of FX mice have higher input resistance and action potential firing rate compared to wild type mice (Domanski et al., 2019). In primary visual cortex, spatial selectivity of cortical FS, PV-positive interneurons is decreased in FX mice (Goel et al., 2018). Many ion channels are altered in a brain region specific manner in FX mice (for review see Deng and Klyachko, 2021)., It is possible that the loss of FMRP has differential effects on FS neurons in the cortex and hippocampus.

### Inhibitory deficits in Fragile X syndrome

There are many types of inhibitory interneurons in the hippocampus that gate activity flow, influence synaptic plasticity, and exhibit control over rhythmic activity (Cobb et al., 1995; Klausberger et al., 2003; Gloveli et al., 2005; Lovett-Barron et al., 2014; Adler et al., 2019). Fast spiking interneurons (FS) make up roughly 14% of the CA1 interneuron population and are located near the pyramidal cell layer or in the stratum oriens where many exhibit strong inhibition on the cell body of pyramidal cells (Bezaire and Soltesz, 2013). FS interneurons in area CA1 provide strong feedforward inhibition as they receive excitatory input from many CA3 pyramidal cells (Buhl et al., 1994; Sik et al., 1995). Low threshold spiking interneurons (LTS) account for approximately 4.5% of total CA1 interneurons and predominantly target the distal dendrites of pyramidal cells in SLM (Bezaire and Soltesz, 2013). LTS interneurons in area CA1 form a canonical feedback inhibition circuit and receive most of their excitatory input from CA1 pyramidal cells (Ali et al., 1998; Klausberger, 2009).

Together FS and LTS inhibitory interneurons play a critical role in balancing excitatory activity (Klausberger and Somogyi, 2008; Pelkey et al., 2017). Proper inhibitory signaling promotes intact plasticity and guides network wide oscillatory events that are critical for learning and memory (Freund and Buzsáki, 1996; Jinno et al., 2007). Several studies on cortical inhibition in FX mice highlighted changes in both FS and LTS interneurons (Gibson et al., 2008; Goel et al., 2018; Domanski et al., 2019). Reduced activation of LTS neurons by metabotropic glutamate receptor signaling and decreased synchronization in synaptic inhibition suggest weakened LTS signaling in somatosensory cortex (Paluszkiewicz et al., 2011). Network hyperexcitability, measured as prolong UP states, in somatosensory cortex is due in part to decreased inhibitory activity from FS neurons (Gibson et al., 2008). These studies demonstrate the importance of investigating specific types of inhibitory interneurons to understand how changes in inhibition contribute to FX pathology.

Although there are several known hippocampal functions that require appropriate stratum oriens inhibitory interneuron signaling (Lovett-Barron et al., 2014; Adler et al., 2019), there are surprisingly few investigations that directly investigate the hippocampal inhibitory interneurons of this region in FX models. While we saw no difference in intrinsic properties of FS neurons between wild type and FX mice, changes in the excitatory synaptic drive onto FS neurons or the inhibitory synaptic drive onto downstream neurons may be different in FX mice. We found that LTH neurons in FX mice were less excitable compared to wild type mice. While the exact axon target in the hippocampus of CA1 LTH neurons remains unknown, the lower intrinsic excitability would reduce inhibitory drive of hippocampal circuits. If like OLM neurons, LTH neurons provide feedback inhibition onto CA1 pyramidal neurons, the reduced excitability of FX LTH neurons would make the synaptic excitation from CA1 neurons less effective, thus reducing feedback inhibition. Prior studies reported reduced inhibitory drive onto pyramidal cells in CA1 in the FX hippocampus (Wahlstrom-Helgren and Klyachko, 2015; Sabanov et al., 2017). It is unclear however, which type of inhibitory interneuron(s) were the source of this inhibition. Additionally, a reduction in inhibitory drive from decreased intrinsic excitability from FX LTH neurons could result in poor modulation of CA1 neuron output. FX mice have been shown to exhibit unstable theta-gamma coupling in the CA1 hippocampus ((Talbot et al., 2018). This discoordination of hippocampal rhythms could be a result of decreased LTH neuron output, and therefore unreliable control, of CA1 pyramidal neuron activity.

FXS patients display several behavioral deficits associated with hippocampal dysfunction such as epilepsy and sensory hypersensitivity. One of the leading hypotheses of these behavioral deficits in FXS is an imbalance of excitatory/inhibitory activity in the hippocampus. Each inhibitory interneuron subtype plays a specific role in orchestrating the flow of excitatory activity through the hippocampus and a deficit in even one subtype in FXS could contribute to hippocampal dysfunction. We found that LTH inhibitory interneurons, a recently identified subclass of low threshold spiking neurons, in the CA1 stratum oriens of the hippocampus in FX mice have higher expression of the hyperpolarization activated cation current, I_h_. The higher I_h_ in FX LTH neurons contributed lower input resistance compared to WT LTH neurons. These changes contribute to lower intrinsic excitability of CA1 stratum oriens LTH neurons in FX mice, which can potentially alter the E/I balance of the CA1 hippocampal circuit. This study highlights the importance of investigating hippocampal interneurons in a subtype and brain region specific fashion which is critical to understanding the cellular basis for neurological deficits in FXS.

## Acknowledgements

We thank Kim Pagtama for assistance with immunohistochemistry, Arsh Ali for the Neurolucida reconstructions, Dr. Richard Gray for essential assistance with analysis and acquisition software, and members of the Johnston and Brager labs for helpful comments and lively discussion of this manuscript. This work was supported by the National Institutes of Health grant R01 MH100510 (DHB), the National Foundation of Sciences GRFP (LTH), and The University of Texas at Austin continuing graduate student fellowship (LTH).

## Author Contributions

L.T.H. and D.H.B. designed experiments; L.T.H. and D.H.B. acquired data; L.T.H. and D.H.B. analyzed data; L.T.H. and D.H.B. interpreted results of experiments; L.T.H. and D.H.B. prepared figures; L.T.H. and D.H.B. drafted manuscript; L.T.H. and D.H.B. edited and revised manuscript.

## Dedication

This manuscript is dedicated to Dr. Richard Gray, our instrumental colleague whose legacy has made a lasting impact to the field of hippocampal neurophysiology.

## References

Adler A, Zhao R, Shin ME, Yasuda R, Gan W-B (2019) Somatostatin-Expressing Interneurons Enable and Maintain Learning-Dependent Sequential Activation of Pyramidal Neurons. Neuron 102:202–216.e7.

Ali AB, Deuchars J, Pawelzik H, Thomson AM (1998) CA1 pyramidal to basket and bistratified cell EPSPs: dual intracellular recordings in rat hippocampal slices. J Physiol 507 (Pt 1):201–217.

Baude A, Bleasdale C, Dalezios Y, Somogyi P, Klausberger T (2007) Immunoreactivity for the GABAA Receptor 1 Subunit, Somatostatin and Connexin36 Distinguishes Axoaxonic, Basket, and Bistratified Interneurons of the Rat Hippocampus. Cerebral Cortex 17:2094–2107.

Bean BP (2007) The action potential in mammalian central neurons. Nat Rev Neurosci 8:451–465.

Bell MV, Hirst MC, Nakahori Y, MacKinnon RN, Roche A, Flint TJ, Jacobs PA, Tommerup N, Tranebjaerg L, Froster-Iskenius U, Kerr B, Turner G, Lindenbaum RH, Winter R, Prembrey M, Thibodeau S, Davies KE (1991) Physical mapping across the fragile X: Hypermethylation and clinical expression of the fragile X syndrome. Cell 64:861–866.

Belmonte MK, Bourgeron T (2006) Fragile X syndrome and autism at the intersection of genetic and neural networks. Nat Neurosci 9:1221–1225.

Bezaire MJ, Soltesz I (2013) Quantitative Assessment of CA1 Local Circuits: Knowledge Base for Interneuron-Pyramidal Cell Connectivity. Hippocampus 23:751–785.

Brager DH, Akhavan AR, Johnston D (2012) Impaired Dendritic Expression and Plasticity of h-Channels in the fmr1−/y Mouse Model of Fragile X Syndrome. Cell Reports 1:225–233.

Brandalise F, Kalmbach BE, Mehta P, Thornton O, Johnston D, Zemelman BV, Brager DH (2020) Fragile X Mental Retardation Protein Bidirectionally Controls Dendritic I _h_ in a Cell Type-Specific Manner between Mouse Hippocampus and Prefrontal Cortex. J Neurosci 40:5327–5340.

Brown MR, Kronengold J, Gazula V-R, Chen Y, Strumbos JG, Sigworth FJ, Navaratnam D, Kaczmarek LK (2010) Fragile X mental retardation protein controls gating of the sodium-activated potassium channel Slack. Nat Neurosci 13:819–821.

Buhl EH, Halasy K, Somogyi P (1994) Diverse sources of hippocampal unitary inhibitory postsynaptic potentials and the number of synaptic release sites. Nature 368:823–828.

Chao H-T, Chen H, Samaco RC, Xue M, Chahrour M, Yoo J, Neul JL, Gong S, Lu H-C, Heintz N, Ekker M, Rubenstein JLR, Noebels JL, Rosenmund C, Zoghbi HY (2010) Dysfunction in GABA signalling mediates autism-like stereotypies and Rett syndrome phenotypes. Nature 468:263–269.

Chittajallu R, Craig MT, McFarland A, Yuan X, Gerfen S, Tricoire L, Erkkila B, Barron SC, Lopez CM, Liang BJ, Jeffries BW, Pelkey KA, McBain CJ (2013) Dual origins of functionally distinct O-LM interneurons revealed by differential 5-HT_3A_R expression. Nature Neuroscience 16:1598–1607.

Cobb SR, Buhl EH, Halasy K, Paulsen O, Somogyi P (1995) Synchronization of neuronal activity in hippocampus by individual GABAergic interneurons. Nature 378:75–78.

Colbert CM, Pan E (2002) Ion channel properties underlying axonal action potential initiation in pyramidal neurons. Nat Neurosci 5:533–538.

Contractor A, Klyachko VA, Portera-Cailliau C (2015) Altered neuronal and circuit excitability in Fragile X Syndrome. Neuron 87:699–715.

DeFelipe J et al. (2013) New insights into the classification and nomenclature of cortical GABAergic interneurons. Nat Rev Neurosci 14:202–216.

Deng P-Y, Carlin D, Oh YM, Myrick LK, Warren ST, Cavalli V, Klyachko VA (2019) Voltage-Independent SK-Channel Dysfunction Causes Neuronal Hyperexcitability in the Hippocampus of Fmr1 Knock-Out Mice. J Neurosci 39:28–43.

Deng P-Y, Klyachko VA (2016) Genetic upregulation of BK channel activity normalizes multiple synaptic and circuit defects in a mouse model of fragile X syndrome. The Journal of Physiology 594:83–97.

Deng P-Y, Rotman Z, Blundon JA, Cho Y, Cui J, Cavalli V, Zakharenko SS, Klyachko VA (2013) FMRP Regulates Neurotransmitter Release and Synaptic Information Transmission by Modulating Action Potential Duration via BK Channels. Neuron 77:696–711.

Domanski APF, Booker SA, Wyllie DJA, Isaac JTR, Kind PC (2019) Cellular and synaptic phenotypes lead to disrupted information processing in Fmr1-KO mouse layer 4 barrel cortex. Nat Commun 10:4814.

Feng Y, Gutekunst CA, Eberhart DE, Yi H, Warren ST, Hersch SM (1997) Fragile X mental retardation protein: nucleocytoplasmic shuttling and association with somatodendritic ribosomes. J Neurosci 17:1539–1547.

Freund TF, Buzsáki G (1996) Interneurons of the hippocampus. Hippocampus 6:347–470.

Fu Y-H, Kuhl DPA, Pizzuti A, Pieretti M, Sutcliffe JS, Richards S, Verkert AJMH, Holden JJA, Fenwick RG, Warren ST, Oostra BA, Nelson DL, Caskey CT (1991) Variation of the CGG repeat at the fragile X site results in genetic instability: Resolution of the Sherman paradox. Cell 67:1047–1058.

Gibson JR, Bartley AF, Hays SA, Huber KM (2008) Imbalance of Neocortical Excitation and Inhibition and Altered UP States Reflect Network Hyperexcitability in the Mouse Model of Fragile X Syndrome. Journal of Neurophysiology 100:2615–2626.

Gloveli T, Dugladze T, Saha S, Monyer H, Heinemann U, Traub RD, Whittington MA, Buhl EH (2005) Differential involvement of oriens/pyramidale interneurones in hippocampal network oscillations in vitro. The Journal of Physiology 562:131–147.

Goel A, Cantu DA, Guilfoyle J, Chaudhari GR, Newadkar A, Todisco B, de Alba D, Kourdougli N, Schmitt LM, Pedapati E, Erickson CA, Portera-Cailliau C (2018) Impaired perceptual learning in a mouse model of Fragile X syndrome is mediated by parvalbumin neuron dysfunction and is reversible. Nat Neurosci 21:1404–1411.

Hagerman RJ, Berry-Kravis E, Kaufmann WE, Ono MY, Tartaglia N, Lachiewicz A, Kronk R, Delahunty C, Hessl D, Visootsak J, Picker J, Gane L, Tranfaglia M (2009) Advances in the Treatment of Fragile X Syndrome. Pediatrics 123:378–390.

Hagerman RJ, Hagerman PJ (2002) The fragile X premutation: into the phenotypic fold. Current Opinion in Genetics & Development 12:278–283.

Hall SS, Lightbody AA, Reiss AL (2008) Compulsive, self-injurious, and autistic behavior in children and adolescents with fragile X syndrome. Am J Ment Retard 113:44–53.

Hewitt LT, Ordemann GJ, Brager DH (2021) High and low expression of the hyperpolarization activated current (Ih) in mouse CA1 stratum oriens interneurons. Physiological Reports 9:e14848.

Hu H, Cavendish JZ, Agmon A (2013) Not all that glitters is gold: off-target recombination in the somatostatin–IRES-Cre mouse line labels a subset of fast-spiking interneurons. Front Neural Circuits 7 Available at: https://www.frontiersin.org/articles/10.3389/fncir.2013.00195/full [Accessed February 25, 2021].

Jinno S, Klausberger T, Marton LF, Dalezios Y, Roberts JDB, Fuentealba P, Bushong EA, Henze D, Buzsáki G, Somogyi P (2007) Neuronal Diversity in GABAergic Long-Range Projections from the Hippocampus. J Neurosci 27:8790–8804.

Kalmbach BE, Johnston D, Brager DH (2015) Cell-Type Specific Channelopathies in the Prefrontal Cortex of the fmr1-/y Mouse Model of Fragile X Syndrome. eNeuro 2 Available at: https://www.eneuro.org/content/2/6/eneuro.0114-15.2015 [Accessed May 17, 2020].

Kaufmann WE, Cortell R, Kau ASM, Bukelis I, Tierney E, Gray RM, Cox C, Capone GT, Stanard P (2004) Autism spectrum disorder in fragile X syndrome: Communication, social interaction, and specific behaviors. American Journal of Medical Genetics Part A 129A:225–234.

Klausberger T (2009) GABAergic interneurons targeting dendrites of pyramidal cells in the CA1 area of the hippocampus. European Journal of Neuroscience 30:947–957.

Klausberger T, Magill PJ, Márton LF, Roberts JDB, Cobden PM, Buzsáki G, Somogyi P (2003) Brain-state- and cell-type-specific firing of hippocampal interneurons in vivo. Nature 421:844–848.

Klausberger T, Somogyi P (2008) Neuronal Diversity and Temporal Dynamics: The Unity of Hippocampal Circuit Operations. Science 321:53–57.

Lien C-C, Martina M, Schultz JH, Ehmke H, Jonas P (2002) Gating, modulation and subunit composition of voltage-gated K ^+^ channels in dendritic inhibitory interneurones of rat hippocampus. The Journal of Physiology 538:405–419.

Lovett-Barron M, Kaifosh P, Kheirbek MA, Danielson N, Zaremba JD, Reardon TR, Turi GF, Hen R, Zemelman BV, Losonczy A (2014) Dendritic Inhibition in the Hippocampus Supports Fear Learning. Science 343:857–863.

Lupica CR, Bell JA, Hoffman AF, Watson PL (2001) Contribution of the Hyperpolarization-Activated Current (*I*_h_) to Membrane Potential and GABA Release in Hippocampal Interneurons. Journal of Neurophysiology 86:261–268.

Maccaferri G, McBain CJ (1996) The hyperpolarization-activated current (Ih) and its contribution to pacemaker activity in rat CAI hippocampal stratum oriens-alveus interneurones. J Physiol:12.

Magee JC (1999) Dendritic Ih normalizes temporal summation in hippocampal CA1 neurons. Nat Neurosci 2:508–514.

Paluszkiewicz SM, Martin BS, Huntsman MM (2011) Fragile X Syndrome: The GABAergic System and Circuit Dysfunction. DNE 33:349–364.

Pelkey KA, Chittajallu R, Craig MT, Tricoire L, Wester JC, McBain CJ (2017) Hippocampal GABAergic Inhibitory Interneurons. Physiological Reviews 97:1619–1747.

Poolos NP, Migliore M, Johnston D (2002) Pharmacological upregulation of h-channels reduces the excitability of pyramidal neuron dendrites. Nat Neurosci 5:767–774.

Routh BN, Johnston D, Brager DH (2013) Loss of Functional A-Type Potassium Channels in the Dendrites of CA1 Pyramidal Neurons from a Mouse Model of Fragile X Syndrome. J Neurosci 33:19442–19450.

Routh BN, Rathour RK, Baumgardner ME, Kalmbach BE, Johnston D, Brager DH (2017) Increased transient Na+ conductance and action potential output in layer 2/3 prefrontal cortex neurons of the fmr1−/y mouse. J Physiol 595:4431–4448.

Rubenstein JLR, Merzenich MM (2003) Model of autism: increased ratio of excitation/inhibition in key neural systems. Genes Brain Behav 2:255–267.

Sabanov V, Braat S, D’Andrea L, Willemsen R, Zeidler S, Rooms L, Bagni C, Kooy RF, Balschun D (2017) Impaired GABAergic inhibition in the hippocampus of Fmr1 knockout mice. Neuropharmacology 116:71–81.

Santoro B, Chen S, Lüthi A, Pavlidis P, Shumyatsky GP, Tibbs GR, Siegelbaum SA (2000) Molecular and Functional Heterogeneity of Hyperpolarization-Activated Pacemaker Channels in the Mouse CNS. J Neurosci 20:5264–5275.

Sik A, Penttonen M, Ylinen A, Buzsaki G (1995) Hippocampal CA1 interneurons: an in vivo intracellular labeling study. J Neurosci 15:6651–6665.

Strumbos JG, Brown MR, Kronengold J, Polley DB, Kaczmarek LK (2010) Fragile X Mental Retardation Protein Is Required for Rapid Experience-Dependent Regulation of the Potassium Channel Kv3.1b. J Neurosci 30:10263–10271.

Talbot ZN, Sparks FT, Dvorak D, Curran BM, Alarcon JM, Fenton AA (2018) Normal CA1 Place Fields but Discoordinated Network Discharge in a Fmr1-Null Mouse Model of Fragile X Syndrome. Neuron 97:684–697.e4.

Tricoire L, Pelkey KA, Erkkila BE, Jeffries BW, Yuan X, McBain CJ (2011) A Blueprint for the Spatiotemporal Origins of Mouse Hippocampal Interneuron Diversity. J Neurosci 31:10948–10970.

Wahlstrom-Helgren S, Klyachko VA (2015) GABAB receptor-mediated feed-forward circuit dysfunction in the mouse model of fragile X syndrome. J Physiol 593:5009–5024.

Zhang Y, Bonnan A, Bony G, Ferezou I, Pietropaolo S, Ginger M, Sans N, Rossier J, Oostra B, LeMasson G, Frick A (2014) Dendritic channelopathies contribute to neocortical and sensory hyperexcitability in Fmr1−/y mice. Nat Neurosci 17:1701–1709.

